# COVID-19 vaccination induces cross-neutralisation of sarbecoviruses related to SARS-CoV-2

**DOI:** 10.1101/2025.04.04.647203

**Authors:** Grace E. West, Rebecca B. Morse, Benjamin L. Sievers, Adam Abdullahi, Kimia Kamellian, Mark Tsz Kin Cheng, The CITIID-NIHR BioResource COVID-19 Collaboration, Douglas G. Stewart, Divya Diamond, David L. Robertson, Spyros Lytras, Sam J. Wilson, Suzannah J. Rihn, Ravindra K. Gupta

## Abstract

Close relatives of SARS-CoV-1 and SARS-CoV-2 continue to circulate in wildlife, posing an ongoing threat of zoonotic spillover. While vaccination played a key role in controlling the COVID-19 pandemic, variants of concern (VOC) with immune evasive substitutions emerged on multiple occasions, causing widespread breakthrough infections. The combined threats of zoonosis and newly emerging VOCs, coupled with the potential for recombination, underscore the need to assess the breadth of existing vaccine-mediated protection. Here, we investigate a cohort of older individuals (median age 68.5 years) for the potential of cross-neutralisation against Omicron lineage VOC and animal sarbecoviruses induced by four COVID-19 vaccine doses. Despite the recent use of a bivalent mRNA vaccine dose (encoding spike from Wu-1 and omicron), we observed that neutralisation of Omicron lineage VOCs such as BA.1 and BA.2 were reduced compared to SARS-CoV-2 Wu-1, suggesting an imprinted immune response from pre-Omicron lineage viruses. Similarly, both SARS-CoV-1 and a SARS-CoV-1-related bat CoV were neutralised less efficiently than SARS-CoV-2 Wu-1. Unexpectedly, however, we observed that two animal SARS-CoV-2-related viruses, BANAL-20-52 (from bats) and a pangolin-CoV, were more sensitive to serum neutralising antibodies than SARS-CoV-2 Wu-1 itself. These surprising findings suggest that vaccine-mediated adaptive immunity may provide efficient cross-protection against certain animal sarbecoviruses.

## Introduction

The importance of zoonotic virus transmission in global public health was highlighted in 2019 when SARS-CoV-2, a coronavirus similar to several circulating coronaviruses in bats, emerged in Wuhan, China (Zhou *et al*., 2020). This marked the third major coronavirus outbreak in humans over the previous two decades, following the SARS and MERS outbreaks of 2003 and 2012. During the COVID-19 pandemic, the swift development of monovalent vaccines based on the spike (receptor binding) protein from the first SARS-CoV-2 sequence isolated from Wuhan in 2019 (SARS-CoV-2 Wu-1) (Zhou *et al*., 2020), was instrumental in controlling transmission and protecting at-risk populations, which included adults aged over 65 and immunocompromised people (Williamson *et al*., 2020).

Despite generally high initial efficacy of these vaccines (Polack *et al*., 2020; Voysey *et al*., 2021), older and immunocompromised individuals had lower neutralising antibody titres (Collier *et al*., 2021; Yang *et al*., 2021) and, across healthy adults, antibody titres were observed to wane more rapidly within six months (Shrotri *et al*., 2021; Haq *et al*., 2024). Thus, booster regimens were introduced; a third dose of monovalent vaccine as a booster established effective neutralising titres against the original Wu-1 variant (Polack *et al*., 2020; Voysey *et al*., 2021), as well as reasonable, short-lived neutralisation against emerging variants of concern (VOC) (Meng *et al*., 2022). VOC are characterised by nucleic acid substitutions throughout the genome (Viana *et al*., 2022), however, there has been specific interest in substitutions in the coronavirus spike, and in particular the receptor binding domain (RBD) that interacts with the host ACE2 receptor. Non-synonymous substitutions within the receptor-binding domain, such as L452R and E484Q, are associated with increased transmissibility, infection and immune escape, which can facilitate breakthrough infections post-vaccination (Cao *et al*., 2022; McCallum *et al*., 2022; Wang *et al*., 2022). Moreover, the emergence of the Omicron lineage VOC dramatically increased infection rates, particularly affecting those aged 65 and above

(Elliott *et al*., 2022; Nyberg *et al*., 2022). Since the recognition of VOC emergence as an ongoing threat to vaccination efforts, bivalent and variant-specific vaccines have been deployed to optimise neutralisation of new VOC that are now recognised as emerging in older and immunosuppressed individuals (Kemp *et al*., 2021; Markov *et al*., 2023).

SARS-CoV-1 and SARS-CoV-2, along with many dozens of other SARS-related coronaviruses (SARSr-CoVs), belong to the *Sarbecovirus* subgenus within the Betacoronavirus genus of the *Coronaviridae* family. The majority of sarbecoviruses have been identified in bats belonging to the *Rhinolophus* genus (horseshoe bats), which are the reservoir for the progenitor viruses of SARS-CoV-1 and SARS-CoV-2 (Zhou *et al*., 2020; Han *et al*., 2023; Latinne *et al*., 2024).

Although sarbecoviruses have been identified in diverse regions globally, most have been sampled from *Rhinolophus* bats in southern China and south east Asia, which often overlap with regions of high human population (Forero-Muñoz *et al*., 2024). This has raised questions about whether other sarbecoviruses from bats or other wild species might have the potential to enter humans and cause future epidemics or pandemics, particularly as previous work has suggested that humans living in these regions are routinely exposed to and infected by other bat SARSr-CoVs (Sánchez *et al*., 2022). Of additional concern is the fact that since SARS-CoV-2 is now endemic in the human population, coinfection of humans has the very real risk of leading to recombination, a common occurrence among bat SARSr-CoVs, which could generate antigenically novel variants (Lytras *et al*., 2022).

There is thus a great interest in identifying both the individual animal sarbecoviruses that may be more likely to enter humans, and what features might enable them to do so. A key determinant of cross-species transmission for animal sarbecoviruses is likely to be the capacity of their spikes to engage angiotensin-converting enzyme 2 (ACE2), the primary human entry receptor for SARS-CoV-1 and SARS-CoV-2, or to utilise human transmembrane protease, serine 2 (TMPRSS2) (Hoffmann *et al*., 2020; Letko, Marzi and Munster, 2020; Zhou *et al*., 2020). Several animal sarbecoviruses have been reported to bind or utilise these human entry pathways (Hoffmann *et al*., 2020; Letko, Marzi and Munster, 2020; Zhou *et al*., 2020; Huang *et al*., 2023), and there is no evidence that any significant adaptation was required prior to spillover (MacLean *et al*., 2021; Havens *et al*., 2025). Moreover, while some bat sarbecoviruses, such as RaTG13 from *Rhinolophus affinis* (Zhou *et al*., 2020), share a relatively high overall genome similarity to SARS-CoV-2, they can have varying levels of similarity in different intragenic regions due to recombination in their ancestry. RaTG13, for example, is more divergent in the region of the viral spike that binds to host receptors (Boni *et al*., 2020), whereas other sarbecoviruses have nearly identical receptor-binding regions. In particular, BANAL-20-52 and BANAL-20-236, which were identified in *Rhinolophus* samples from northern Laos, differ from SARS-CoV-2 by only one or two amino acid residues in the receptor-binding motif (Temmam *et al*., 2022). In addition, some sarbecoviruses isolated from pangolins have spikes which share close genetic similarity to the spike of SARS-CoV-2, and appear to be able to enter human cells, raising questions as to whether pangolins could be conceivable intermediate hosts in the spillover of SARS-CoV-2 from bats to humans (Xiao *et al*., 2020; Nga *et al*., 2022; Hou *et al*., 2023).

Given that some animal sarbecoviruses appear poised to potentially spillover into humans once more, it is crucial to understand whether existing COVID-19 vaccine-mediated immunity might offer protection against these sarbecoviruses. This may be particularly important for individuals at higher risk of severe disease and will also enable the development of forward-thinking vaccination strategies. In this study of older adults in the UK at higher risk of severe COVID-19, we investigate whether COVID-19 vaccines offer cross-protection against Omicron VOC and SARS-related bat and pangolin sarbecoviruses that can utilise ACE2, some of which have fewer substitutions within the receptor-binding regions of their spikes than VOC relative to SARS-CoV-2 Wu-1. We aim to improve understanding of population preparedness for potential sarbecovirus spillover events and examine the role existing vaccines may have in future epidemic or pandemic mitigation.

## Results

### Neutralisation sensitivity of BA.2-derived VOC following a fourth vaccine dose in older adults

To first investigate existing immunity in older individuals to Omicron variants of concern (VOC) compared to the original SARS-CoV-2 Wu-1 variant, we utilised an age- and sex-matched cohort (total n = 22, median age = 68.5 years) of individuals in the United Kingdom who received a bivalent SARS-CoV-2 (Wu-1/BA.1) vaccine as their fourth dose in late 2022. These individuals had previously received a primary two-dose series of either AZD1222 (adenovirus vector-based) or BNT162b2 (mRNA-based) 3 months apart followed by a third mRNA-based monovalent booster dose 6 months after the primary series **(Figure 1A, Table 1)**. We performed a luciferase-based neutralisation assay in HeLa cells exogenously expressing human ACE2 using serum taken ~30 days (median: 31.5, IQR: 29-33) after the fourth bivalent vaccine dose and assessed neutralisation capacity against SARS-CoV-2 Wu-1 and a range of VOC from the Omicron lineage **(Figure 1B**). We assessed neutralisation against previously circulating variants (BA.2 and its descendants BA.4 and BA.5, which share identical spike sequences) and descendants of BA.2 that had not yet circulated at the time of vaccination (BA.2.86 and XBB). As expected, the highest levels of neutralisation were observed against SARS-CoV-2 Wu-1 (geometric mean titre (GMT) = 54410) **(Figure 1C, Supplementary Figure 1A**), despite the vaccine including BA.1, suggesting an imprinted immune response towards the Wu-1 original variant to which these individuals were first exposed. Despite significantly lower neutralisation of BA.2 and BA.4/5 compared to SARS-CoV-2 Wu-1, neutralising titres remained high (GMT = 10084 and 7312, respectively). Four doses did not induce cross-protection against BA.2.86 and XBB variants (GMT = 399 and 343, respectively), which circulated later, reflecting significant evasion of vaccine-induced immunity due to antigenic divergence of these spikes.

**Table 1.**
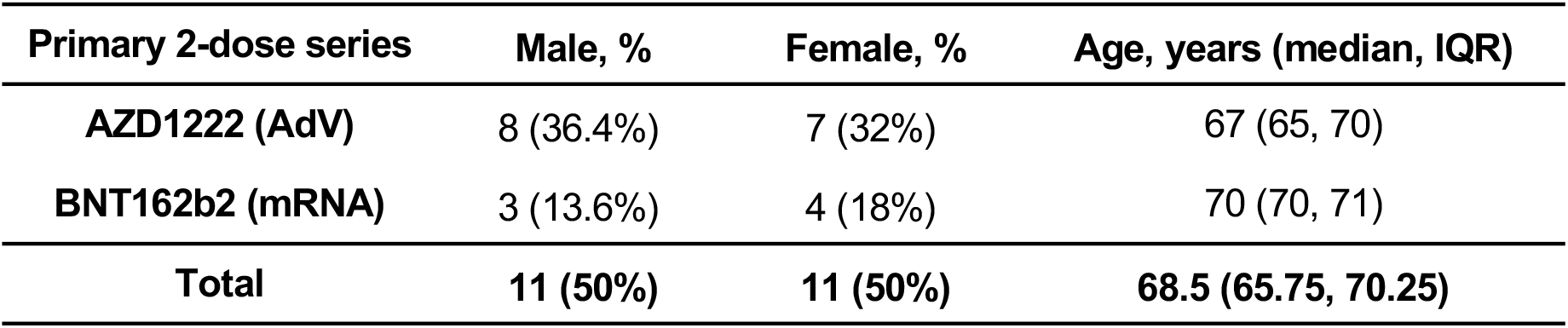
Cohort characteristics. Age- and sex-matched individuals (n = 22) vaccinated four times against SARS-CoV-2. All individuals received a primary two-dose series of either AZD1222 (adenovirus vector-based) or BNT162b2 (mRNA-based) against SARS-CoV-2 Wuhan-Hu-1 (Wu-1), followed by a 3rd dose (mRNA-based) against Wu-1 and a 4th bivalent dose (mRNA-based) against Wu-1 and B.1.1.529 (Omicron BA.1).

**Figure 1.**
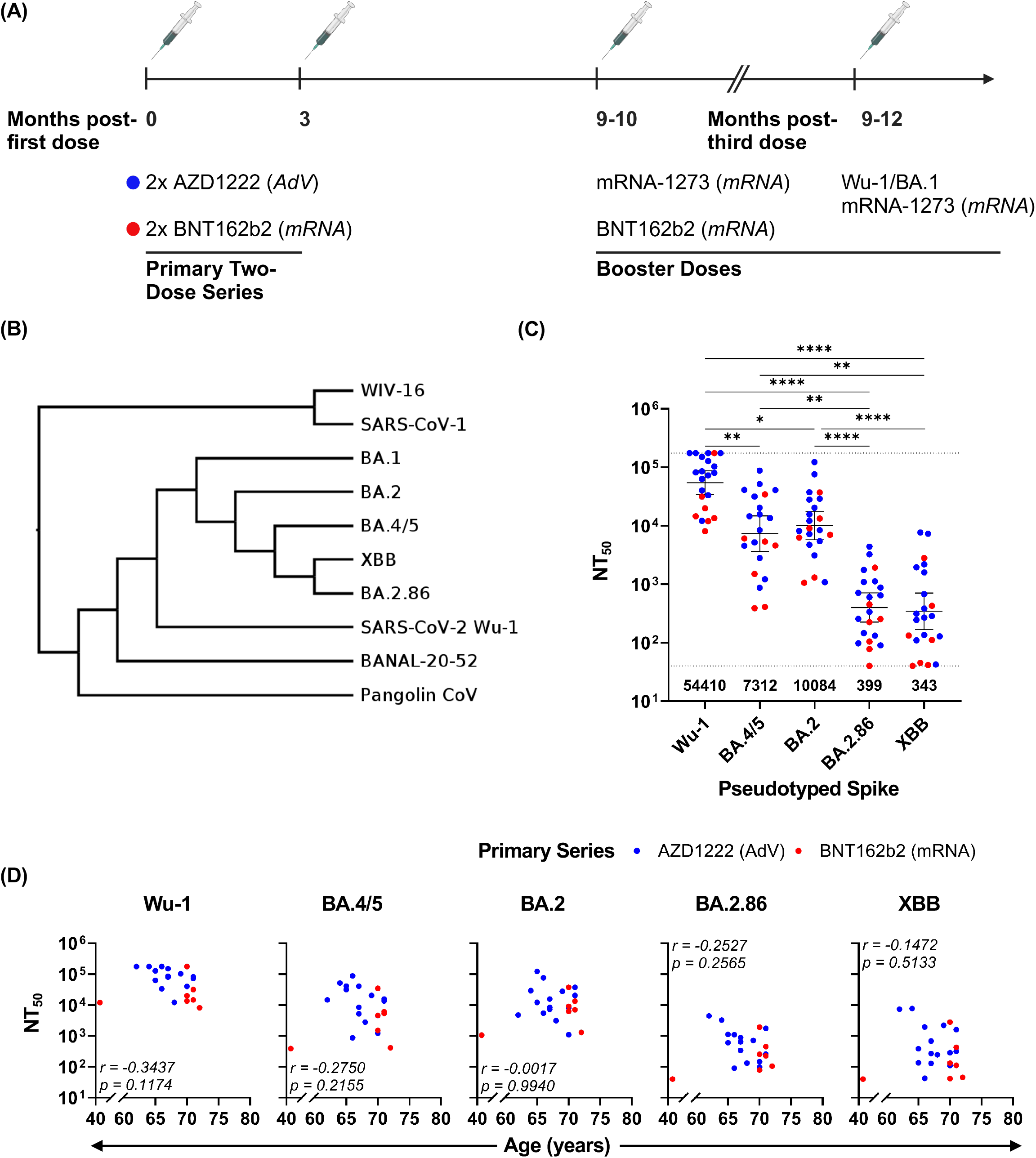
Multi-dose vaccine schedules in a cohort of older individuals with mixed infection histories induce strong humoral responses against SARS-CoV-2 Wu-1 and early Omicron lineages. (A) Schematic of cohort vaccination schedule for first four doses and vaccines received. AdV = Adenovirus vector-based vaccine, mRNA = mRNA-based vaccine. Created in BioRender. Morse, R. (2025) https://BioRender.com/p69s364. (B) Nucleotide sequence-based spike protein phylogenetic cladogram of SARS-CoV-2 Wu-1, SARS-CoV-2 variants and related sarbecoviruses. (C) Luciferase neutralisation assay of SARS-CoV-2 spike-pseudotyped lentiviruses after preincubation with n = 22 serum samples taken 1 month post-fourth dose in HeLa cells stably expressing human ACE2. 50% neutralising titres (NT50) of cohort sera against spike-pseudotyped lentiviruses are shown with individual points (blue = primary two-dose series with AZD1222, red = primary two-dose series with BNT162b2) and bars indicating geometric mean titre (GMT) with 95% CI. Dotted lines indicate the minimum and maximum detection limits of the neutralisation assay. GMT is written below the minimum detection limit dotted line. P-values were calculated using the Friedman test and Dunn’s Multiple Comparisons test. (D) Correlation of age and NT50 against spike-pseudotyped lentiviruses. Spearman r values and p-values were calculated using the nonparametric Spearman correlation test. *p < 0.05; **p < 0.01; ***p < 0.001; ****p < 0.0001; ns, p > 0.05.

Above age 60, there was no correlation between age and NT50 (the half-maximal inhibitory concentration) in our cohort (**Figure 1D**). Only neutralisation against Wu-1 was significantly different between primary series groups, although neutralising titres remained high for both groups (**Supplementary Figure 1B**). There was a trend of older age in individuals who had received the BNT162b2 primary series (**Supplementary Figure 1C**), which was due to the prioritisation of older individuals for the first available vaccines in the UK (Materials and Methods: Cohort and Ethical Approval).

### Diverse sarbecoviruses can use either human or R. sinicus ACE2 for cell entry

We next wanted to investigate what level of cross-protection this vaccine-mediated immunity might offer against animal sarbecoviruses. To do this, we identified a panel of 40 different sarbecoviruses from bats, humans and pangolins for which full-length genome sequences were available and generated a phylogenetic tree to illustrate the relationships between these viruses **(Figure 2A, Supplementary Table 1)**. Due to high levels of recombination between sarbecoviruses, different regions of the genome can have strikingly different phylogenetic relationships, so for this analysis, the tree was inferred using the nucleotide sequence corresponding to the receptor binding domain of each spike.

**Figure 2:**
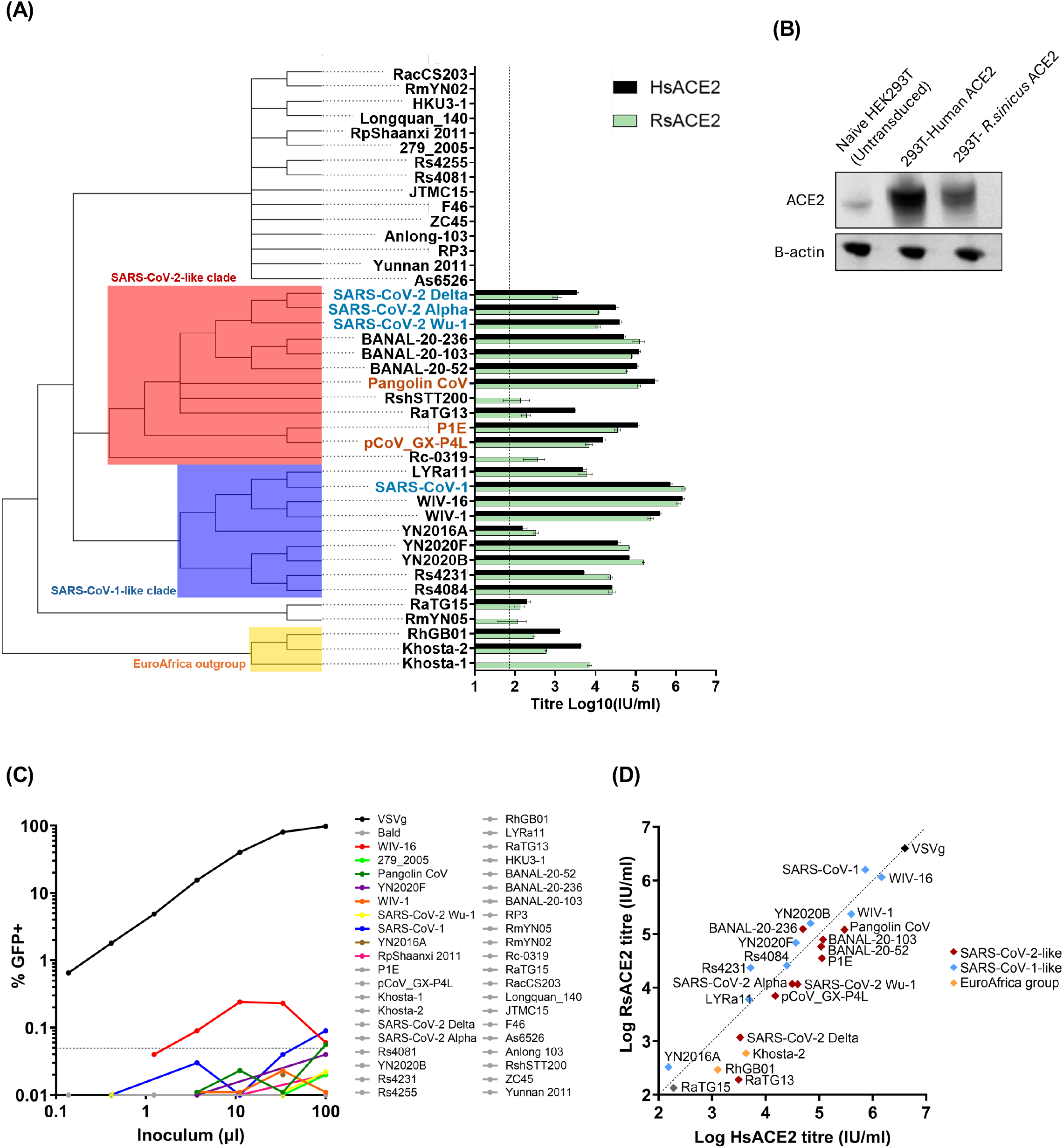
Entry of sarbecovirus spikes using human and *Rhinolophus sinicus* ACE2. (A) Nucleotide sequence-based phylogeny of the receptor-binding domains in sarbecovirus spikes isolated from bats (black text), pangolins (red text) and humans (blue text). Nodes with ultrafast bootstrap support below 80% have been dropped. Different clades are highlighted on the cladogram: Red = SARS-CoV-2-like clade, blue = SARS-CoV-1-like clade, yellow = EuroAfrica outgroup. Mean viral titres of sarbecovirus spike-pseudotyped lentiviruses in HEK293T cells stably expressing ACE2 from humans (black bars) or *Rhinolophus sinicus* (green bars). Dotted line represents minimum threshold for detection. (B) Expression of ACE2 detected in untransduced HEK293T cells vs HEK293T cells transduced with human or *Rhinolphus sinicus* ACE2 by western blot. B-actin = loading control. (C) Entry of sarbecovirus spike-pseudotyped lentiviruses into untransduced 293T cells (Positive control = VSVg). Dotted line represents minimum threshold for detection. (D) Comparison of usage of human and *Rhinolophus sinicus* ACE2 by different sarbecovirus spikes. Blue = SARS-CoV-1-like clade, red = SARS-CoV-2-like clade, yellow = EuroAfrica Outgroup. Dotted line represents equal use of human and *Rhinolophus sinicus* ACE2.

The spikes clustered into distinct clades, which have been previously described based on sequence and receptor binding (Cui, Li and Shi, 2019; Letko, Marzi and Munster, 2020; Guo *et al*., 2021). The two main clades contained spikes that were either closely related to SARS-CoV-1 or those which were more closely related to SARS-CoV-2. Three spikes were in the EuroAfrica clade, which contains viruses which have been isolated from both Europe and Africa, though the three spikes from our panel were all isolated from European bats.(Crook *et al*., 2021; Seifert *et al*., 2022). There were also a number of other outliers which did not cluster with the above three clades and these spikes are known to contain deletions which may affect their capacity to use ACE2 (Hu *et al*., 2017; Wells *et al*., 2021).

The ability for the spikes included in the phylogenetic analysis above to use ACE2 from either humans or *Rhinolophus sinicus* (a species believed to have hosted SARS-CoV-2 progenitor viruses (Lytras *et al*., 2022), and which hosts many SARSr-CoVs, **Supplementary Table 1**) to enter human cells was then assessed using an entry assay utilising sarbecovirus spike-pseudotyped particles. Briefly, lentivirus particles encoding a GFP reporter (Bainbridge *et al*., 2001) were pseudotyped with the individual sarbecovirus spikes, and entry of the pseudotyped particles into naïve 293T cells, or 293T cells exogenously expressing human or *Rhinolophus sinicus* ACE2 was then quantified using flow cytometry to measure GFP positive cells.

To confirm that any entry we observed in our assay was due to exogenous ACE2 expression, we first measured entry of the sarbecovirus spike-pseudotyped particles into naïve 293T cells which express very low levels of endogenous ACE2 **(Figure 2B)**. In line with previous findings, the sarbecovirus spike-pseudotyped particles were generally unable to enter these cells **(Figure 2C)**, although spikes from SARS-CoV, WIV-16 and pangolin-CoV facilitated a very low level of entry close to the detection threshold, likely reflecting use of a small amount of endogenous ACE2, or an ACE2-independent pathway. Nevertheless, since all three barely exceeded our approximate limit for detection (~0.05% GFP+ cells), we can conclude that sarbecovirus spikes are generally unable to enter naïve 293T cells.

We then examined the level of entry the sarbecovirus-pseudotyped particles displayed in 293T cells exogenously expressing human or *Rhinolophus sinicus* ACE2 and found that many spikes from the SARS-CoV-1-like and SARS-CoV-2-like clades efficiently used human ACE2 for entry, often with a comparable efficiency to ACE2 from *Rhinolophus sinicus* **(Figure 2A, D)**.

Interestingly, spikes from the SARS-CoV-1-like clade often had marginally higher entry titres in cells expressing *Rhinolophus sinicus* ACE2, while spikes from the SARS-CoV-2-like clade typically had higher entry titres in the cells expressing human ACE2 (**Figure 2D**). Moreover, several bat and pangolin sarbecoviruses from the SARS-CoV-2-like clade facilitated higher levels of entry than any of the SARS-CoV-2 variants that we tested. While spikes from the EuroAfrica group exhibited some entry in the presence of exogenous ACE2, this was typically lower and more species-specific than spikes more closely related to SARS-CoV-1 or SARS-CoV-2. Other spikes which were more distantly related to SARS-CoV-1 and SARS-CoV-2 failed to use human or *Rhinolophus sinicus* ACE2 **(Figure 2A)**. These spikes may be able to use ACE2 orthologues from other *Rhinolophus* species, or they may use an unknown, ACE2-independent entry pathway (Guo *et al*., 2022; Khaledian *et al*., 2022).

### Four COVID-19 vaccine doses in a population with mixed infection history are associated with very high neutralisation of sarbecoviruses related to SARS-CoV-2

Given that a number of the circulating bat and pangolin sarbecovirus spikes we tested appeared able to efficiently use human ACE2 for entry, and therefore could be candidates for potential spillover events, we next sought to investigate whether vaccines for SARS-CoV-2 could provide cross-protection against entry of a selection of these sarbecoviruses. BANAL-20-52 and pangolin-CoV were selected as close relatives of SARS-CoV-2 which can efficiently use human ACE2 for entry **(Figure 1B, 2A)**. The spike receptor binding motifs (RBM) of BANAL-20-52 and pangolin-CoV are highly similar to the RBM of the SARS-CoV-2 Wu-1 spike, only varying at residue 498, while their RBDs differ by only six or seven amino acids, respectively **(Figure 3A)**. We also selected SARS-CoV-1 and a close bat sarbecovirus relative, WIV-16, because they both used human ACE2 effectively for entry **(Figure 2A)** and we were interested in determining whether any potential vaccine-mediated cross protection might extend to SARS-CoV-1-like sarbecoviruses. SARS-CoV-1 and WIV-16 accordingly have more divergent spike sequences, with 75.82% and 76.84% amino acid similarity compared to SARS-CoV-2, respectively.

**Figure 3:**
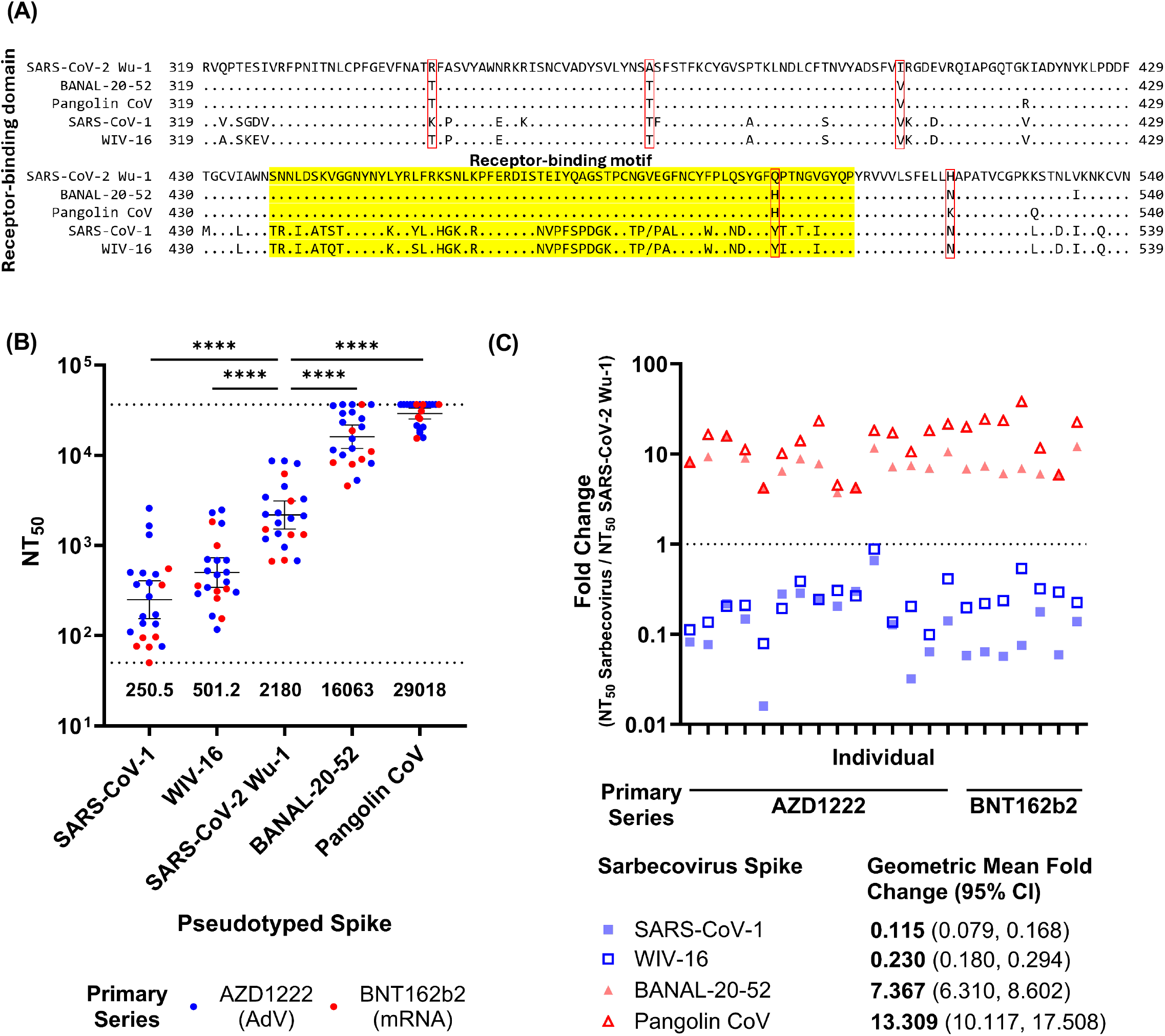
Neutralisation of sarbecovirus spikes by sera from quadruple-vaccinated individuals. (A) Alignment of the amino acid sequence of the receptor-binding domains of the spikes in this experiment. Yellow = Receptor-binding motif. Amino acid residues different between BANAL-20-52/Pangolin CoV and SARS-CoV-2 are outlined in red. (B) GFP neutralisation assay of sarbecovirus spike-pseudotyped lentiviruses after preincubation with n = 22 serum samples taken 1 month post-fourth dose in HEK293T cells stably expressing human ACE2. 50% neutralising titres (NT50) of cohort sera against spike-pseudotyped lentiviruses are shown with individual points (blue = primary two-dose series with AZD1222, red = primary two-dose series with BNT162b2) and bars indicating geometric mean titre (GMT) with 95% CI. Dotted lines indicate the minimum and maximum detection limits of the neutralisation assay. GMT is written below the minimum detection limit dotted line. Representative graph of n = 2 independent experiments. P-values were calculated using the Wilcoxon matched-pairs signed rank test relative to Wu-1. *p < 0.05; **p < 0.01; ***p < 0.001; ****p < 0.0001; ns, p > 0.05. (C) Fold change in neutralisation of lentiviruses pseudotyped with sarbecovirus spikes compared with SARS-CoV-2 Wu-1 (dotted line at y = 1) for each individual. Geometric mean fold change and 95% CI were calculated to compare neutralisation of each pseudotyped sarbecovirus spike to SARS-CoV-2 Wu-1. Representative graph of n = 2 independent experiments.

To investigate neutralisation, we incubated sera with sarbecovirus spike-pseudotyped particles before assessing entry into HEK293T cells expressing exogenous human ACE2. We found that SARS-CoV-1 and WIV-16 were both poorly neutralised compared to SARS-CoV-2 Wu-1 (GMT = 250.4 and 501.2, respectively, compared to 2180) **(Figure 3B, Supplementary Figure 2A**), as might be predicted from their genetic distance from SARS-CoV-2. Surprisingly, BANAL-20-52 and pangolin-CoV were neutralised significantly better than SARS-CoV-2 Wu-1, with NT50 values approximately 7.4-fold and 13.3-fold higher than that of SARS-CoV-2 Wu-1 (GMT = 16063 and 29018, respectively, compared to 2180), and with similar neutralisation breadth across all participants **(Figure 3B, C, Supplementary Figure 2A)**. We did not observe an impact of primary series or age on neutralisation of each sarbecovirus **(Supplementary Figure 2B, C)**.

## Discussion

In this study we investigated whether neutralising antibody responses elicited by four doses of COVID-19 vaccines in older patients could offer cross-protection against SARS-CoV-2 VOC and other bat and pangolin sarbecoviruses that are able to readily use ACE2 to enter human cells.

Compared to the original SARS-CoV-2 Wu-1 variant, we observed markedly reduced cross-neutralisation of Omicron VOC, consistent with the evolution of their spike proteins in response to population level selective pressure from vaccines and prior infection. Notably, we found that bivalent vaccination induced significant protection against BA.2 and BA.4/5, which was expected given that BA.2 and BA.1 differ by only 6 amino acids in their RBD, coupled with the fact that these variants circulated prior to vaccination, so it is possible that participants could have been exposed before samples were collected or had cross-protection from BA.1 exposure. In contrast, our tested sera exhibited minimal neutralisation of emerging variants BA.2.86 and XBB, consistent with their antigenic difference from Wu-1 and emergence after the administration of the fourth vaccine dose, where prior exposure to XBB and BA.2.86 was unlikely. Similarly, particles pseudotyped with SARS-CoV-1 or a SARS-CoV-1-like spike (from bat sarbecovirus WIV-16) exhibited little or reduced neutralisation compared to SARS-CoV-2 Wu-1. We were therefore quite surprised to observe that individuals receiving a bivalent (Wu-1/BA.1) fourth dose generated neutralising titres over 7-fold higher against BANAL-20-52 and pangolin-CoV as compared to SARS-CoV-2 Wu-1.

One reason this finding was particularly unexpected is that BANAL-20-52 and pangolin-CoV exhibit only one amino acid difference from SARS-CoV-2 Wu-1 in the receptor binding motif (at position 498). Although substitutions in the receptor-binding motif (RBM) are known to be significant determinants of ACE2 binding affinity and immune evasion (Cao *et al*., 2022; McCallum *et al*., 2022), the magnitude of change in neutralisation between these CoVs and SARS-CoV-2 Wu-1 was unanticipated. Whereas increased neutralisation of RaTG13 has been observed relative to SARS-CoV-2, likely because of its decreased affinity for human ACE2 (Wrobel *et al*., 2020; Cantoni *et al*., 2022), the receptor-binding domains of pangolin-CoV and BANAL-20-52 have both been observed to bind the human ACE2 with similar, or higher, efficiency (Niu *et al*., 2021; Temmam *et al*., 2022). Differences in neutralisation may be due to other changes in the RBD, or even in other parts of the spike.

Indeed, it is plausible that structural differences such as altered RBD exposure or spike trimer configuration, potentially modulated by allosteric interactions across the spike (from NTD to S2) play a critical role in the pattern of neutralisation profiles that we observe in our current study (Wrobel *et al*., 2021; Ou *et al*., 2023). Future studies employing domain-swapping of sarbecovirus spikes, or with structural studies such as cryo-electron microscopy, could reveal more about which regions or conformations of sarbecovirus spikes influence neutralisation. Our findings highlight the importance of understanding cross-neutralisation among sarbecoviruses that utilise ACE2, with implications for pan-sarbecovirus vaccine design and zoonotic spillover preparedness.

Interestingly, previous studies have demonstrated that after three vaccine doses, neutralising antibodies show greater tolerance to substitutions of the RBD and there is expansion of B cell subsets with broader reactivity due to somatic hypermutation (Andreano *et al*., 2023). In individuals aged over 70, subsequent vaccine doses have been shown to further refine memory B cell differentiation, favouring atypical memory B cell subsets which can produce broadly neutralising antibodies (Ferreira *et al*., 2023). To this end, other previous work has already shown that after a booster vaccination, there were higher neutralising titres against SARS-CoV-2 Wu-1 as well as omicron–boosted cross-reactivity (Garcia-Beltran *et al*., 2022). This phenomenon could offer one possible explanation as to why we observed such high levels of cross-protection against BANAL-20-52 and pangolin-CoV, which possess conserved RBD sequences. This raises the possibility that vaccination could be useful in preventing spillover of some emerging sarbecoviruses in specific regions where there is a higher chance of contact with infected wildlife. The addition of the Omicron-adapted final vaccine dose in our cohort could also have contributed to improved cross-neutralisation, as structural and antigenic differences in the spikes from Wu-1 and Omicron variants might elicit an immune memory which is more tolerant of differences in the spike sequence. This highlights a potential for tailored booster strategies with the aim of eliciting a more broadly reactive immune memory. This has important implications for the design of next generation sarbecovirus vaccines including the incorporation of conserved spike epitopes into multivalent or mosaic formulations spanning pre-emergent and emergent lineages, providing broader protection against both circulating VOCs and zoonotic sarbecoviruses with pandemic potential.

However, it is also worth noting that our sarbecovirus neutralisation results differ from some published reports, particularly with respect to pangolin-CoV. While we observed significantly better neutralisation of particles pseudotyped with the spike from pangolin-CoV than with the SARS-CoV-2 Wu-1 spike, other studies reported comparable or lower neutralisation (Tan Chee-Wah *et al*., 2021; Hou *et al*., 2023; Huang *et al*., 2023). Many factors including prior infection, hybrid immunity and methodological variability could explain these differences. One key consideration is the number of vaccine doses that participants had received; the aforementioned studies used sera from participants after their first or second dose, whilst we tested sera from participants who had received four vaccine doses and were conceivably exposed to and/or infected by one or more VOC. As mentioned above, breadth of neutralisation appears to increase with additional vaccine doses (Andreano *et al*., 2023; Ferreira *et al*., 2023), which could explain the improved cross-neutralisation of pangolin-CoV after four doses. In accordance, Zhang et al. reported that the neutralisation of two different pangolin sarbecoviruses (one of which was closely related to pangolin-CoV used in this study) increased with subsequent vaccine doses (Zhang *et al*., 2024), although neutralisation in their study was reported as a percentage at a fixed serum dilution of 1:25 rather than as NT50, which limits the comparison of neutralisation of SARS-CoV-2 Wu-1 and the pangolin CoVs since they were all completely neutralised at this serum dilution in our study. By contrast, we chose to titrate serum samples over a broad range to calculate NT50 values that should better capture the nuances of neutralisation. This was particularly important since many participants’ sera completely neutralised some of the sarbecovirus spike-pseudotyped particles at our starting dilution of 1:50.

Our study has a few limitations, including the relatively small cohort size and the unknown SARS-CoV-2 infection history of our cohort participants. However, the widespread infection rates in this age group in the UK (Elliott *et al*., 2022) indicate a high likelihood that many in the cohort had hybrid immunity, particularly with Omicron lineage VOC infections. Additionally, the consistency in breadth of cross-neutralisation across all participants despite differing primary series (**Figure 3C**) mitigates some concerns about NT50 variability and the effects of infection on cross-protection. Additionally, the availability of cohort serum samples restricted the number of sarbecovirus spikes that we could test for neutralisation; larger scale investigations including spikes from more diverse sarbecoviruses with varied human ACE2 binding capabilities would provide a more detailed picture of differences in neutralisation across the sarbecovirus subgenus.

Ultimately, we believe our study underscores the importance of understanding vaccine-induced cross-neutralisation for future epidemic and pandemic preparedness. The enhanced neutralisation of BANAL-20-52 and pangolin-CoV observed here suggests that existing vaccines, particularly with repeated boosters, may provide a foundation for broader protection against other animal sarbecoviruses. Further research should investigate the basis for increased neutralisation sensitivity of BANAL-20-52 and pangolin-CoV spike that we observed, including exploring the role of other spike domains or certain spike conformations, as this may be key to identifying the determinants of sarbecovirus sensitivity to neutralisation and unlocking improvements to future vaccines. Our findings highlight the value of booster vaccines in driving cross-sarbecovirus neutralisation, positioning current vaccines as a potential first line of defence against future zoonotic sarbecovirus outbreaks.

## Acknowledgments

This work was funded by a Wellcome Senior Fellowship to RKG (WT108082AIA), a UKRI Horizon Europe Guarantee (Frontier Research Guarantee) award to SJR (EP/Y011414/1 for an ERC Starting Grant) and Medical Research Council (https://www.ukri.org/councils/mrc) awards MR/P022642/1 (SJW, SJR), MR/V01157X/1 (SJW, SJR) and MC UU 00034/5 (DLR). GEW is funded by the NIHR Cambridge Biomedical Research Centre. RBM is funded by the Harding Distinguished Postgraduate Scholars Programme. AA was supported by the Cambridge-Africa award and Harvard Takemi Program in International Health. This research was also supported by the NIHR Cambridge Biomedical Research Centre (NIHR203312). The SARS CoV-2 antibody responses in immunocompromised patients study is funded by Addenbrooke’s Charitable Trust (ACT) and Vasculitis UK. The views expressed are those of the authors and not necessarily those of the NIHR or the Department of Health and Social Care.

## Materials and methods

### Cohort and ethical approval

We selected 22 serum samples from participants who had received four vaccine doses. The cohort median age was 68.5 years (**Table 1**), and selected participants were sex- and age-matched. For the primary two-dose series, 7 individuals received BNT162b2 mRNA-based vaccines and 15 received AZD1222 adenovirus vector-based vaccines. Participants were recruited through the NBR118 Study at the Cambridge NIHR BioResource Centre. Individuals consented to providing post-vaccination biological samples as well as relevant clinical data. The study was approved by the East of England – Cambridge Central Research Ethics Committee (17/EE/0025) on April 28th, 2020.

In the UK, the vaccination response began with the licensing of the mRNA-based Pfizer vaccine (BNT162b2), initially offered to residents of care homes and individuals over 80 years old.

Subsequently the adenovirus vector-based Oxford-AstraZeneca ChAdOx1 nCoV-19 vaccine (AZD1222) was offered to the next priority group. Despite the initial success, neutralising antibody responses were observed to wane over time, leading to the recommendation of booster doses. Many individuals aged over 70 have now received at least four doses, with ongoing vaccinations enhancing both the potency and breadth of the neutralising antibody response against SARS-CoV-2 (Andreano *et al*., 2023).

### Phylogeny and alignment

We identified a number of sarbecoviruses for which the full genome sequence was available **(Supplementary Table 1)**. For these sarbecoviruses, the nucleotide sequence of the RBD regions was aligned using MAFFT (localpair option) (Katoh and Standley, 2013), and the tree was inferred using IQ-TREE2 under a GTR+F+R10 substitution model with 1000 ultrafast bootstrap iterations to determine node support (Minh *et al*., 2020). The resulting tree was rooted by the European viruses and nodes with confidence below 80% were dropped. The cladogram was visualised using RStudio.

Amino acid sequences of sarbecovirus spike proteins were aligned using MAFFT v7.526, implemented via Python 3.8.8 with Biopython 1.83 for sequence processing (Cock *et al*., 2009). The custom Python script is available at https://github.com/baseten418/shirley.

### Cell maintenance

HEK293T cells, a human kidney cell line, were used in experiments and for generation of cell lines and were obtained from existing stocks from the Rihn laboratory. All HEK293T derived cell lines were cultured in Dulbecco’s Modified Eagle Medium (DMEM) supplemented with 10% fetal bovine serum (FBS) and 50 mg/ml gentamicin and grown at 37°C in 5% CO2. Cells were routinely screened for mycoplasma.

HeLa-ACE2 cells (a gift from J. Voss) were maintained in DMEM supplemented with 10% FBS and 1% penicillin-streptomycin and grown at 37C in 5% CO2.

### Generation of ACE2-expressing cell lines

HEK293T cell lines exogenously expressing human (HsACE2) (NP_001358344.1) or *Rhinolophus sinicus* (RsACE2) (XP_019601896.1) ACE2 were generated via lentivirus transduction using the pLV-EF1a-IRES-hygro expression vector (Addgene #85134). Plasmid sequences were verified by Plasmidsaurus using Oxford Nanopore Technology with custom analysis and annotation.

Lentiviruses were produced by co-transfecting HEK293T cells with 5μg of the ACE2 expression vector, 5μg of a lentiviral transfer and packaging plasmid (NL4.3 GagPol), and 1μg of VSVg (glycoprotein from vesicular stomatitis virus) using polyethylenimine (PEI) in serum-free DMEM. Cells were seeded at 1/3 confluence 24 hours prior to transfection. Viral supernatant was collected at 48 hours post-transfection and filtered (0.45µm), and the filtered supernatant was then used to transduce HEK293T cells in 24-well plates by spinoculation (1 hour, 1600 RPM, room temperature), after which cells were incubated at 37°C in 5% CO2. The transduced cells were subsequently selected and maintained in 200µg/ml hygromycin for at least seven days before use, and mock-transduced controls confirmed successful selection.

### Western blot analyses

To prepare cell lysates, cell pellets were resuspended in protein sample buffer (12.5% glycerol, 175LmM Tris-HCl [pH 8.5], 2.5% SDS, 70□mM 2-mercaptoethanol, 0.5% bromophenol blue). Proteins were separated on 4% to 12% Bis-Tris polyacrylamide gels and transferred onto nitrocellulose membranes. Blots were probed with either anti-ACE2 (21115-1-AP; Proteintech) or anti-β-actin (66009-1-1g; Proteintech), then were probed with fluorescently labelled goat antimouse (SA5-10176; ThermoFisher) or goat anti-rabbit (SA5-10036; ThermoFisher) secondary antibodies and scanned using a LiCor Odyssey scanner.

### Generation of luciferase-expressing VOC spike-pseudotyped lentiviruses

Pseudotyped particles representing the original SARS-CoV-2 variant (Wu-1) (MN908947.3) and VOC were generated as previously described (Kamelian *et al*., 2025). In short, specific amino acid substitutions were introduced into the D614G pCNA_SARS-CoV-2_S plasmids. The pseudotyped particles were created using a triple plasmid transfection system, where the Spike-expressing plasmid, along with a lentiviral packaging vector (p8.91), and a luciferase expression vector (psCSFLW), were transfected into HEK293T cells using the Fugene HD transfection reagent (Promega). After 48 hours, the viruses were collected and stored at –80C. Individual TCID50s were determined by titrating the viruses on HeLa cells expressing exogenous human ACE2.

### Neutralisation assay of luciferase-expressing VOC spike-pseudotyped lentiviruses

Neutralisation assays were conducted using HeLa cells expressing exogenous human ACE2 and SARS-CoV-2 spike-pseudotyped particles that carry the luciferase gene. The pseudotyped particles were incubated with serially diluted heat-inactivated sera from vaccinated individuals, in duplicate for 1 hour at 37C. Controls for cells only and virus with cells (i.e., no serum) were included on each plate. After the incubation, HeLa-ACE2 cells were added to each well. After 48 hours at 37C and 5% CO2, luminescence was measured using the BrightGlo Luciferase Assay System (Promega, UK), and neutralisation was calculated relative to the controls. The 50% neutralization titer (NT50) was determined using the half-maximal inhibitory concentration values of individual serum samples, normalized to control infections, from their serial dilutions, and reported as geometric mean titres. Statistical comparisons among groups were performed using the Friedman test and Dunn’s Multiple Comparisons test. All statistical analyses were conducted on Prism version 10.4.1.

### Generation of sarbecovirus spike-pseudotyped lentiviruses

Lentiviruses pseudotyped with sarbecovirus spikes were produced by co-transfection of 5μg of CSGW reporter lentiviral vector that encodes GFP (Bainbridge *et al*., 2001), 5 μg of NL4.3 GagPol, and 8 μg of the sarbecovirus spike (codon optimised, in expression vector pcDNA3.1). After 48h, the sarbecovirus spike-pseudotyped particles were harvested and passed through 0.45 um filters. Accession IDs for each sarbecovirus spike protein investigated are listed in **Supplementary Table 1**.

### Entry/neutralisation assays of sarbecovirus spike-pseudotyped lentiviruses

24 hours before transduction, 293T-HsACE2 (hs, *Homo sapiens*) or 293T-RsACE2 (rs, *Rhinolophus sinicus*) cells were seeded into 96-well plates at 15,000 cells per well.

For entry assays, sarbecovirus spike-pseudotyped particles were titrated onto cells using 7-point, three-fold serial dilutions in DMEM. Transduced cells were spinoculated for 1 hour at 1600 RPM and 20°C, then incubated for 48 hours at 37°C in 5% CO2. Infected cells were fixed in 4% PFA and stored at 2-5°C for subsequent flow cytometry analysis.

For neutralisation assays, heat-inactivated human serum samples at a starting dilution of 1:50 were titrated using seven point, three-fold serial dilutions in DMEM, then mixed with sarbecovirus spike-pseudotyped particles (which were titrated beforehand to determine a fixed concentration which would infect 5-10% of 293T-HsACE2 cells), followed by incubation at 37° C and 5% CO2 for 1h. Each plate included cell-only and virus- and cell-only (i.e., no serum) controls. Then, the incubated serum/pseudotyped particle mixtures were added to the pre-seeded 293T-HsACE2 cells and spinoculated for 1 hour at 1600 RPM and 20°C, then incubated for 48 hours at 37°C in 5% CO2. Infected cells were fixed in 4% PFA and stored at 2-5°C for flow cytometry analysis.

### Flow cytometry and other analyses

Flow cytometry was conducted using a Cytek® Guava® easyCyte™ Flow Cytometer to measure GFP signal. Neutralisation was calculated relative to cells-only and virus-only (without serum) controls. NT50 values for each group were reported as geometric mean titres, and statistical comparisons between SARS-CoV-2 Wu-1 and the other groups were performed using the Wilcoxon matched-pairs signed rank test. All statistical analyses were conducted on Prism version 10.4.1.

