## supplementary material for "COVID-19 vaccination induces cross-neutralisation of sarbecoviruses related to SARS-CoV-2"

| Spike Protein | Accession ID | Host Species | Location | Date |
|---|---|---|---|---|
| SARS CoV-1 (HsZcc) | AY394995 | Homo sapiens | Guangdong, China | 2003 |
| SARS-CoV-2 Wu-1 (D614G) | MN908947.3 | Homo sapiens | Wuhan, China | 2019 |
| SARS-CoV-2 Alpha | EPI_ISL_3980577 | Homo sapiens | Britain | 2021 |
| SARS-CoV-2 Delta | EPI_ISL_1635330 | Homo sapiens | India | 2021 |
| SARS-CoV-2 Omicron BA.2 | UJP23605.1 | Homo sapiens | USA | 2022 |
| SARS-CoV-2 Omicron BA.4 | UPP14409.1 | Homo sapiens | USA | 2022 |
| SARS-CoV-2 Omicron BA.2.86 | WOY09184.1 | Homo sapiens | South Africa | 2023 |
| SARS-CoV-2 Omicron XBB | OP607807.1 | Homo sapiens | USA | 2022 |
| P1E | EPI_ISL_410539 | Manis javanica | Guanxi, China | 2017 |
| Pangolin CoV | EPI_ISL_410721 | Manis javanica | Guangdong, China | 2019 |
| pCoV_GX-P4L | MT040333.1 | Manis javanica | Guanxi, China | 2017 |
| RhGB01 | MW719567.1 | Rhinolophus hipposideros | Britain | 2020 |
| RShSTT200 | EPI_ISL_852605 | Rhinolophus shameli | Cambodia | 2010 |
| BANAL-20-103 | MZ937001.1 | Rhinolophus pusillus | Fueng, Laos | 2020 |
| BANAL-20-236 | MZ937003.1 | Rhinolophus marshalli | Fueng, Laos | 2020 |
| BANAL-20-52 | MZ937000.1 | Rhinolophus malayanus | Fueng, Laos | 2020 |
| Khosta-1 | MZ190137.1 | Rhinolophus ferrumequinum | Russia | 2022 |
| Khosta-2 | MZ190138.1 | Rhinolophus hipposideros | Russia | 2022 |
| RP3 | DQ071615 | Rhinolophus pearsonii | Guanxi, China | 2004 |
| Anlong-103 | KY770858 | Rhinolophus sinicus | Guizhou, China | 2013 |
| Longquan_140 | KF294457 | Rhinolophus monoceros | Guizhou, China | 2012 |
| HKU3-1 | DQ022305 | Rhinolophus sinicus | Hong Kong | 2005 |
| 279_2005 | DQ648857 | Rhinolophus macrotis | Hubei, China | 2004 |
| Rc-o319 | LC556375 | Rhinolophus cornutus | Japan | 2013 |
| RpShaanxi 2011 | JX993987 | Rhinolophus pusillus | Shaanxi, China | 2011 |
| RacCS203 | MW251308 | Rhinolophus acuminatus | Thailand | 2020 |
| As6526 | KY417142 | Aselliscus stoliczkanus | Yunnan, China | 2014 |
| F46 | KU973692 | Rhinolophus pusillus | Yunnan, China | 2012 |
| JTMC15 | KU182964 | Rhinolophus ferrumequinum | Yunnan, China | 2013 |
| LYRa11 | KF569996 | Rhinolophus affinis | Yunnan, China | 2011 |
| RaTG13 | EPI_ISL_402131 | Rhinolophus affinis | Yunnan, China | 2013 |
| RaTG15 | GWHBAUP01000001 | Rhinolophus affinis | Yunnan, China | 2015 |
| RmYN02 | EPI_ISL_412977 | Rhinolophus malayanus | Yunnan, China | 2019 |
| RmYN05 | MZ081376.1 | Rhinolophus malayanus | Yunnan, China | 2020 |
| Rs4081 | KY417143 | Rhinolophus sinicus | Yunnan, China | 2012 |
| Rs4084 | KY417144 | Rhinolophus sinicus | Yunnan, China | 2012 |
| Rs4231 | KY417146.1 | Rhinolophus sinicus | Yunnan, China | 2013 |
| Rs4255 | KY417142 | Rhinolophus sinicus | Yunnan, China | 2013 |
| WIV-1 | KF367457.1 | Rhinolophus sinicus | Yunnan, China | 2012 |
| WIV-16 | KT444582.1 | Rhinolophus sinicus | Yunnan, China | 2013 |
| YN2016A | OK017847.1 | Rhinolophus sinicus | Yunnan, China | 2016 |
| YN2020B | OK017852.1 | Rhinolophus sinicus | Yunnan, China | 2020 |
| YN2020F | OK017856.1 | Rhinolophus sinicus | Yunnan, China | 2020 |
| Yunnan 2011 | JX993988 | Chaerephon plicata | Yunnan, China | 2011 |
| ZC45 | MG772933 | Rhinolophus sinicus | Zheijang, China | 2017 |

**Supplementary Table 1: Accession IDs, host species and location/year of sampling for each sarbecovirus investigated in this study**

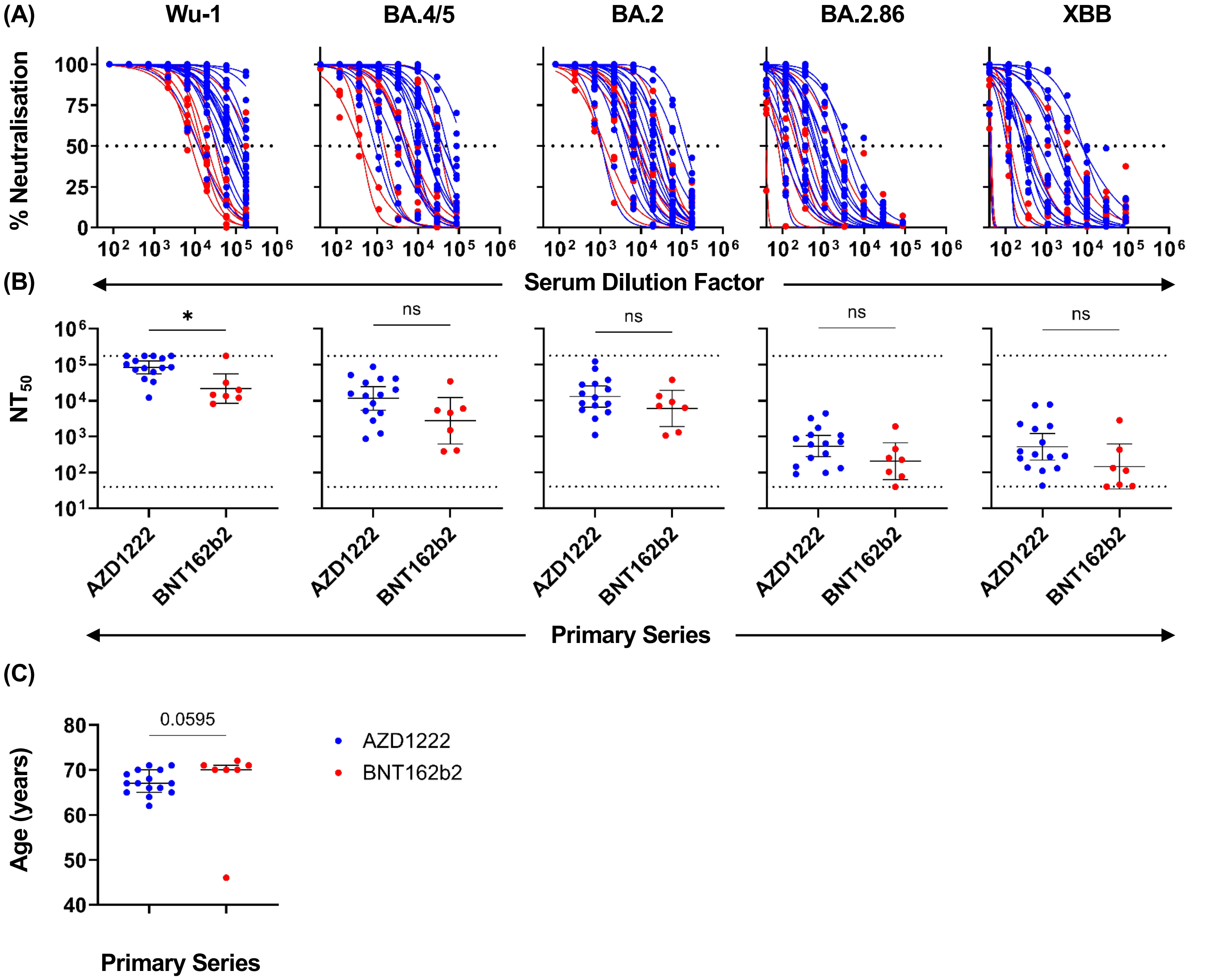


**Supplementary Figure 1: Neutralisation curves and comparisons among replicates, sex, age, and NT50 values using a luciferase readout.** (A) Neutralisation of SARS-CoV-2 spike-pseudotyped luciferase lentiviruses (Wu-1, BA.4/5, BA.2, BA.2.86, XBB) after preincubation with n = 22 serum samples taken 1 month post-fourth dose in HeLa cells stably expressing human ACE2. Values were normalised to cells-only and virus-only wells. Blue points indicate a primary two-dose series with AZD1222 and red points indicate a primary two-dose series with BNT162b2. The dotted line at y = 50 indicates the point at which 50% of the pseudotyped virus was neutralised by serum sample. (B) 50% neutralising titre (NT50) stratified by primary vaccine series (AZD1222 or BNT162b2) with bars indicating geometric mean titre (GMT) with 95% CI. P-values were calculated using a Mann Whitney test. (C) Ages of individuals in the cohort stratified by primary vaccine series. Bars indicate median age with interquartile range. *p < 0.05; **p < 0.01; ***p < 0.001; ****p < 0.0001; ns, p > 0.05.

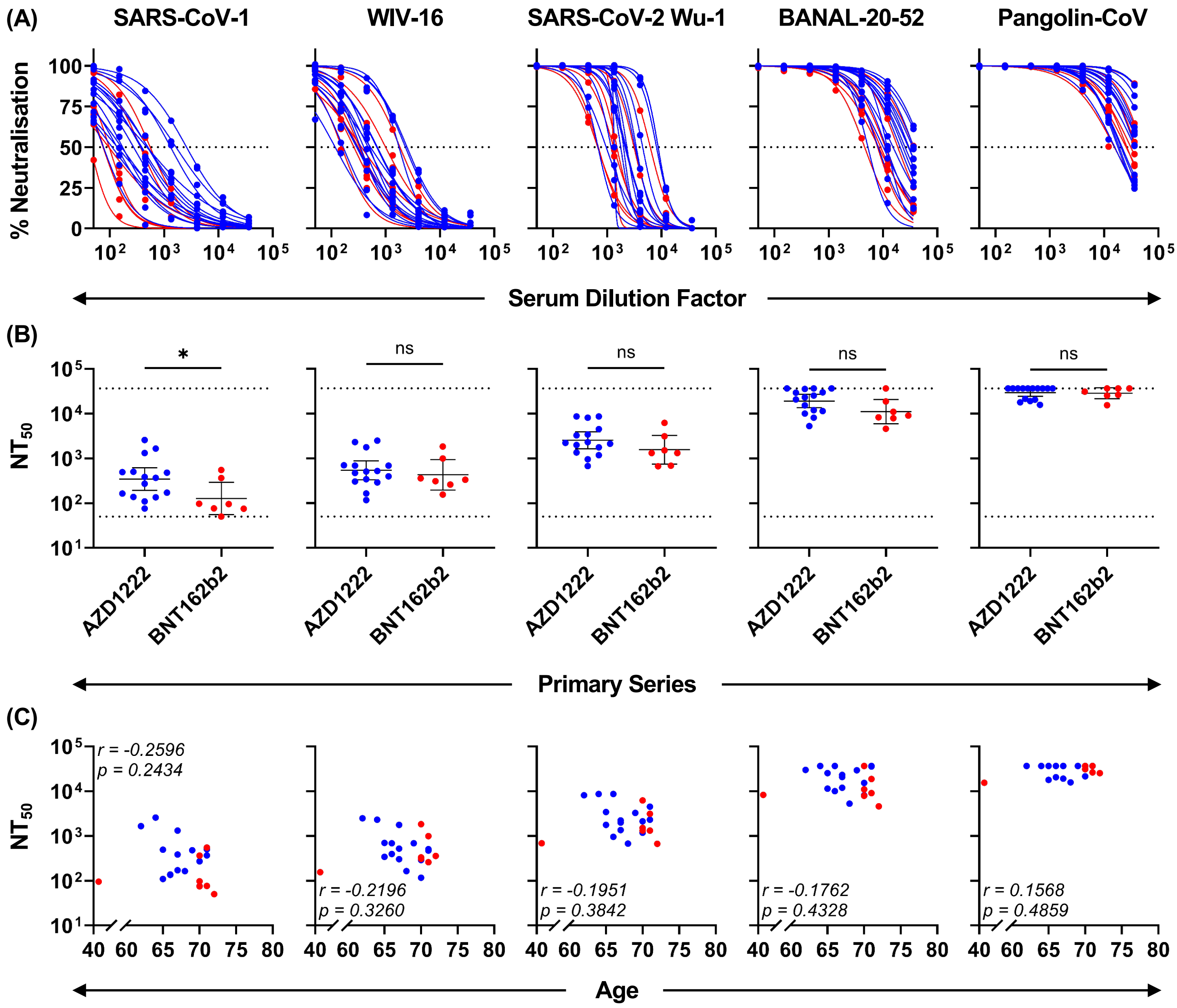


**Supplementary Figure 2: Neutralisation curves and comparisons among replicates, sex, age, and NT50 values using a GFP readout.** (A) Neutralisation of sarbecovirus (SARS-CoV-1, WIV16, BANAL-20-52, Pangolin CoV) and SARS-CoV-2 Wu-1 spike-pseudotyped GFP lentiviruses after preincubation with n = 22 serum samples taken 1 month post-fourth dose in HEK293T cells stably expressing human ACE2. Values were normalised to cells-only and virus-only wells. Blue points indicate a primary two-dose series with AZD1222 and red points indicate a primary two-dose series with BNT162b2. The dotted line at y = 50 indicates the point at which 50% of the pseudotyped virus was neutralised by each serum sample. (B) 50% neutralising titres (NT50) stratified by primary vaccine series (AZD1222 or BNT162b2) with bars indicating geometric mean titre (GMT) with 95% CI. P-values were calculated using a Mann Whitney test. (C) Correlation of age and NT50 against spike-pseudotyped lentiviruses. Spearman r values and p-values were calculated using the nonparametric Spearman correlation test. *p < 0.05; **p < 0.01; ***p < 0.001; ****p < 0.0001; ns, p > 0.05.
